# An efficient error correction and accurate assembly tool for noisy long reads

**DOI:** 10.1101/2023.03.09.531669

**Authors:** Jiang Hu, Zhuo Wang, Zongyi Sun, Benxia Hu, Adeola Oluwakemi Ayoola, Fan Liang, Jingjing Li, José R. Sandoval, David N. Cooper, Kai Ye, Jue Ruan, Chuan-Le Xiao, De-Peng Wang, Dong-Dong Wu, Sheng Wang

## Abstract

Long read sequencing data, particularly those derived from the Oxford Nanopore (ONT) sequencing platform, tend to exhibit a high error rate. Here, we present NextDenovo, a highly efficient error correction and assembly tool for noisy long reads, which achieves a high level of accuracy in genome assembly. NextDenovo can rapidly correct reads; these corrected reads contain fewer errors than other comparable tools and are characterized by fewer chimeric alignments. We applied NextDenovo to the assembly of high quality reference genomes of 35 diverse humans from across the world using ONT Nanopore long read sequencing data. Based on these *de novo* genome assemblies, we were able to identify the landscape of segmental duplications and gene copy number variation in the modern human population. The use of the NextDenovo program should pave the way for population-scale long-read assembly, thereby facilitating the construction of human pan-genomes, using Nanopore long read sequencing data.

## Introduction

An accurate and complete genome is a prerequisite for research into the evolution of species. Third-generation long-read sequencing platforms, such as PacBio single-molecule real-time (SMRT)^1^ and Oxford Nanopore (ONT)^2^, promise to overcome the challenges that are inherent to short-read sequencing and have the potential to resolve most complex and repetitive genomic regions. To this end, they have become the mainstream method of sequencing for genome assembly. The high-fidelity (HiFi) reads recently produced by PacBio display superior performance to *de novo* assembly^3–5^. However, they usually have an average length of ~15 kilobases (kb), and hence are unable to span long tandem or multi-copy highly homologous repeats, which occur widely throughout large genomes, but very specifically in some regions such as centromeres^3, 6^. ONT sequencing can generate >100-kb “ultra-long” reads, which can be used to fill the final gaps of an assembly, most of which are located in these regions^7, 8^. This approach was first used successfully in the assembly of a human centromere (chromosome Y,^9^) and an entire chromosome (chromosome X,^10^), and then combined with HiFi data to assemble a complete human genome^8^. Despite these successes, a single linear reference genome is insufficient to represent the entire genome sequence of a species, and there is an urgent need to construct pan-genomes for population genome research^11–13^. ONT sequencing is characterized by lower cost, higher throughput and a faster turnaround time than PacBio HiFi sequencing and, since it requires less genomic DNA, it can be used anywhere for sampling and sequencing by portable devices. It is therefore eminently suitable for pan-genome projects, especially those with limited budgets or urgent deadlines.

For genome assembly from noisy long ONT reads, two commonly used strategies have been employed, viz. “correction then assembly” (CTA) and “assembly then correction” (ATC); the former (such as Necat^14^ and Canu^15^) are usually slower than the latter (such as Wtdbg2^16^ and Flye^17^), because read-level error correction requires rather more computational resources than contig-level polishing. However, in terms of the assembly of segmental duplications/repeats, and especially for large plant genome assemblies, the CTA-based strategy usually has enhanced ability to distinguish different gene copies and produce more accurate and continuous assemblies.

Here, we present NextDenovo, a highly efficient error correction and CTA-based assembly tool for noisy long reads. We first provide an overview of the NextDenovo pipeline, and then compare it to other error correction and assembly tools using 4 non-human genomes and 35 human genomes. We show that NextDenovo represents an optimal choice for error-correction and genome assembly when working with noisy long reads, especially for large repeat-rich genomes.

## Results

### Overview of the NextDenovo pipeline

As with other CTA assemblers, NextDenovo first detects the overlapping reads (**Fig. 1a**), then filters out the alignments caused by repeats and finally splits the chimeric seeds based on the overlapping depth (**Fig. 1b**). NextDenovo employs the kmer score chain (KSC) algorithm which was used by our previously published polisher tool, NextPolish^18^, to perform the initial rough correction (**Fig. 1c**). Because a large number of noisy or incorrect overlap alignments exist in regions with high error rates, these regions tend to be located within repeat regions. These regions are usually characterized by lower accuracy after the initial correction, but they are nevertheless important for distinguishing different repeat copies during the subsequent graph cleaning procedure. So NextDenovo used a heuristic algorithm to detect these low score regions (LSRs) during the traceback procedure within the KSC algorithm. For the LSRs, a more accurate algorithm, derived by combining the partial order alignment (POA)^19^ and KSC, was used. In detail, each subsequence spanning an LSR was collected, and a kmer set at the flanking sequences of this LSR was generated. Then each subsequence was assigned a matched kmer score based on this kmer set. Subsequences with a lower kmer score (usually caused by heterozygosity or repeats) were filtered out. The longest six subsequences ranked by kmer score were used to produce a pseudo-LSR seed by a greedy POA consensus algorithm. All pseudo-LSR seeds from the same seed were linked as the reference, and all subsequences from this seed were mapped to this reference and the KSC algorithm applied again to produce a corrected pseudo seed. This procedure was called multiple times in order to improve the accuracy of the LSRs. Finally, each LSR was extracted from the corrected pseudo seed and inserted into the corresponding position of the primary corrected seed as the final corrected seeds (**Fig. 1d**).

**Fig. 1:**
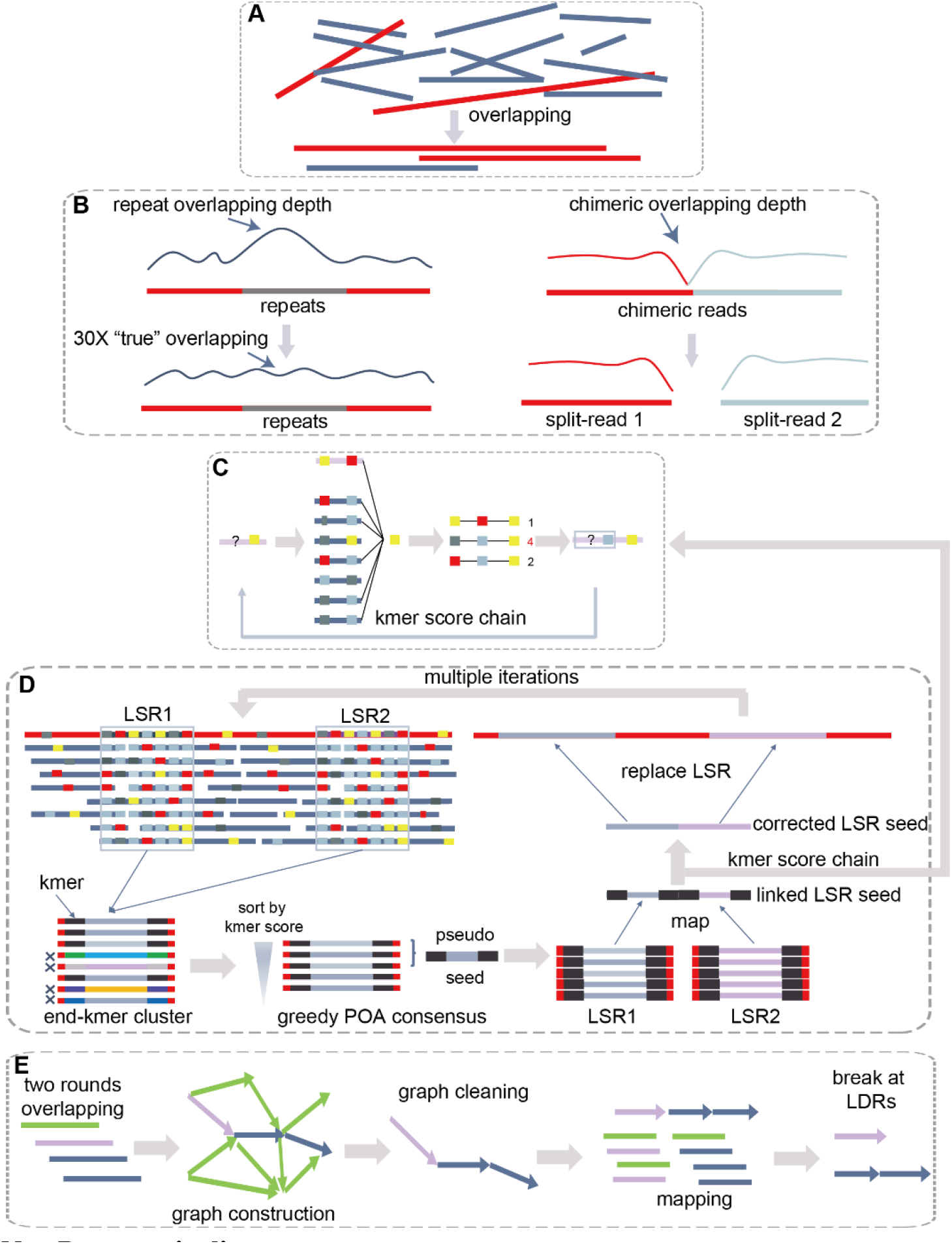
NextDenovo pipeline. (**A**) Overlapping reads. (**B**) Alignments erroneously caused by repeats were filtered out and chimeric reads were split. (**C**) A confidence score was calculated for a given allele at each position with a fixed 3-mer, and the allele with the maximum score was selected as the correct base. The colored rectangles represent the different bases. (**D**) NextDenovo first identities all LSRs at the raw reads, extracts each subsequence spanning these LSRs and assigns a kmer score to each subsequence. Subsequently, NextDenovo filters out the subsequences with lower scores and produces a pseudo-LSR seed using a greedy POA consensus algorithm, all pseudo-LSR seeds from the same seed being linked as the reference, and all subsequences being mapped to this reference whilst the KSC algorithm is reapplied in order to produce a corrected pseudo seed. Then, the corrected LSRs are inserted into the corresponding positions in the raw reads to generate the final corrected reads. (**E**) NextDenovo calculates dovetail alignments by two rounds of overlapping, constructs an assembly graph, removes transitive edges, tips, bubbles and edges with low scores, and generates contigs. Finally, NextDenovo maps all seeds to contigs and breaks a contig if it possesses low quality regions.

The corrected seeds were subjected to two rounds of pairwise overlapping to identify dovetail alignments. The first round used an efficacious parameter set designed to rapidly detect candidate dovetail alignments, which usually contain incorrect alignments or imprecise alignment boundaries. So, for these candidate dovetail alignments, a strict parameter set was used to produce more accurate alignments. Next, a directed string graph was constructed and transitive edges were removed as with most existing assemblers. We used the “best overlap graph” (BOG) algorithm to remove edges for non-repeat nodes (repeat nodes were defined as nodes with indegree or outdegree larger than a thread). For repeat nodes, we found that the BOG algorithm usually removes the corrected edges and breaks the graph connectivity. To fix this problem, we only removed a repeat edge if its alignment identity, length and transitive score (see “Methods”) were less than their corresponding threads. Subsequently, tips were removed and bubbles were resolved. Finally, the graph usually contained some linear paths (no branches and repeated nodes) connecting some complex subgraphs that contained many repeat nodes. We used a greedy progressive graph cleaning strategy to simplify these complex subgraphs. That is, a series of increasingly stringent thresholds were used to filter edges while maintaining connectivity between incoming and outgoing nodes. Finally, all paths were broken at the node connecting with multi-paths, and contigs were outputted from these broken linear paths. To further reduce the possibility of misassemblies, we mapped all seeds to the contigs and broke a contig at the connection point between two nodes if it had a lower mapping depth region (LDR, **Fig. 1e**).

### Benchmarking the error correction module

We benchmarked the error correction performance of NextDenovo against Canu (v2.0) and Necat (v0.0.1) using simulated data and actual biological data based on chr.1 of a human genome (CHM13, **Table 1, Supplementary Table 1**)^8^. In terms of the correction speed, NextDenovo was 7.44, 1.13 times faster on simulated data and 69.25, 1.63 times faster on actual biological data than Canu and Necat, respectively. The differences between simulated data and actual biological data arise because the actual biological data are ONT “ultra-long” reads, the reads to be corrected having an average length of 91.21 kb, 3.99 times longer than the simulated data. This implies that the correction speed advantage of NextDenovo will become increasingly obvious as the read length increases. For the corrected data size, NextDenovo can correct 2.21%, 4.54% more data than Canu, but 1.65%, 1.00% less data than Necat, on simulated data and actual biological data, respectively. Importantly, the average error rate of corrected reads by NextDenovo is 1.82%, 1.31% lower than Canu, and 0.35%, 0.09% lower than Necat, on simulated data and actual biological data, respectively. The average accuracy of corrected reads by NextDenovo is higher than >99%, which is close to the accuracy of PacBio HiFi reads, whereas the corrected reads have a much longer length than HiFi reads. Moreover, the consistent error rate of corrected reads is important for subsequent graph cleaning procedures, because reads alignment identity can be used to distinguish ambiguous edges in the assembly graph, especially for edges from different repeat copies. We found that the ONT reads from the actual biological data usually contain LSRs, and benefit from the heuristic algorithm used by NextDenovo to correct the LSRs with multiple iterations, producing ~89.31% corrected reads that have ≥97% accuracy, whilst only 80.64% for Canu, 88.67% for Necat and 0.18% for the raw data. This is also the reason why some reads are not split or trimmed at the LSRs, resulting in the reads corrected by NextDenovo having a longer average length (90.98 kb) than those corrected by Canu (84.27 kb) and Necat (89.17 kb). The chimeric reads are usually detrimental to assembly graph construction, leading to misconnections and incorrect assembly results. NextDenovo can detect these chimeric reads and can split them at the LSRs; 89.10% of the corrected reads can be mapped to reference with ≥99% coverage, compared to 85.13% for Canu, whilst the comparable figure for Necat was slightly lower (88.30%) than with NextDenovo. This result is consistent with the corrected reads by NextDenovo having the fewest chimeric alignments. In summary, NextDenovo can correct reads at a faster speed; these corrected reads contain fewer errors and are characterized by a more uniform error rate and fewer chimeric alignments.

**Table 1:** Statistics of ONT read error correction.

| Source | Software | Corrected bases rate (%) | Average length (bp) | Max length (bp) | Reads with chimeric alignments (%) | Mapped with $\geq 99\%$ coverage (%) | Mapped with $\geq 97\%$ identity (%) | Average error rate (%) | Wall clock time (hour) |
| --- | --- | --- | --- | --- | --- | --- | --- | --- | --- |
| Simulation (chr1, 62X) | Raw Reads | - | 22,884 | 279,538 | 0.03 | 73.28 | 0.00 | 12.37 | - |
|  | NextDenovo | 83.47 | 23,515 | 188,302 | 0.03 | 99.58 | 99.96 | 0.20 | 2.43 |
|  | Necat | 85.12 | 23,542 | 264,959 | 0.08 | 98.46 | 98.83 | 0.55 | 2.75 |
|  | Canu | 81.26 | 23,369 | 279,129 | 0.41 | 98.03 | 96.58 | 2.02 | 18.08 |
| CHM13 (chr1, 72X) | Raw Reads | - | 91,209 | 499,238 | 17.07 | 81.19 | 0.18 | 8.57 | - |
|  | NextDenovo | 97.13 | 90,981 | 505,469 | 10.70 | 89.10 | 89.31 | 0.90 | 1.83 |
|  | Necat | 98.13 | 89,170 | 506,817 | 11.43 | 88.30 | 88.67 | 0.99 | 2.98 |
|  | Canu | 92.59 | 84,270 | 502,469 | 13.44 | 85.13 | 80.64 | 2.21 | 126.72 |

Only the primary alignments defined by minimap2 of each read were used for evaluation. Corrected base rate is the ratio of the size of the corrected reads to the size of the raw reads to be corrected. Reads with chimeric alignments are defined as reads that have supplementary alignments. Average error rate only uses the reads that are mapped with $\geq 80\%$ coverage. All the software was tested on the same computer with 32 CPUs and 252 GB RAM of memory.

### Assembly evaluation on non-human genomes

We first evaluated NextDenovo in the context of the assembly of four non-human genomes (*Arabidopsis thaliana, Drosophila melanogaster, Oryza sativa* and *Zea mays*) in relation to the most widely used assemblers, Necat (v0.0.1), Canu (v2.0), Flye (v2.8) and Wtdbg2 (v2.5) on ONT data (**Supplementary Table 1**), and then used QUAST (v5.2.0)^20^ to evaluate all assemblies with regard to completeness (assembly size, gene completeness), accuracy (number of misassemblies and phred-scaled base error rate (QV)) and continuity (NG50/LG50 and NGA50/LGA50, **Table 2, Supplementary Table 2**). For the *A. thaliana* and *D. melanogaster* genomes, since the structure of these two genomes is relatively simple, most assemblers produced good assemblies. NextDenovo, Necat and Flye performed better than Canu and Wtdbg2 on the overall evaluation metric, whilst NextDenovo, Necat and Flye reported similar values for completeness and continuity. With regard to accuracy, compared with Necat and Flye, the NextDenovo assemblies contained fewer misassemblies and had a higher QV on the *D. melanogaster* genomes, but the NextDenovo assembly contained 2 more misassemblies and showed a slightly smaller QV than the Flye assembly on the *A. thaliana* genome. The *O. sativa* and *Z. mays* genomes contained more repeats and are more complex. Benefiting from the high-accuracy data after error correction, NextDenovo can more reliably distinguish different repeats, ensuring that the NextDenovo assemblies exhibit greater continuity than other assembler results, especially for the *Z. mays* genome. NextDenovo can deliver an assembly with about 2, 61, 15, 758 times the NGA50 values of Necat, Canu, Flye and Wtdbg2, respectively. Moreover, the NextDenovo assemblies also contained the smallest number of misassemblies and had a higher QV than the other tools. With regard to completeness, the assemblies produced by NextDenovo, Necat, Canu and Flye exhibit similar values in terms of assembly size and gene completeness. Indeed, for the *A. thaliana* and the *O. sativa* genomes, NextDenovo provided near-chromosome-level assemblies because the LGA90 values of these two assemblies were only 10 and 20, implying that most of the chromosomes in these two genomes contain only 1-2 long contigs.

**Table 2:** Statistics of nonhuman assemblies.

| Sample | Software | Assembly size (Mb) | NG50 (Mb)/LG50 | NGA50 (Mb)/LGA50 | No. of misassemblies | QV | Gene completeness (%) | Wall clock time (hour) |
| --- | --- | --- | --- | --- | --- | --- | --- | --- |
| *A. thaliana* (452X) | NextDenovo | 128.37 | 15.18/5 | 15.18/5 | 19 | 33.25 | 99.20 | 6.83 |
|  | Necat | 124.55 | 15.01/5 | 14.98/5 | 44 | 31.93 | 99.20 | 6.82 |
|  | Canu | 138.29 | 9.31/5 | 9.31/6 | 430 | 25.09 | 99.20 | 312.13 |
|  | Flye | 121.16 | 14.63/5 | 14.63/5 | 17 | 35.65 | 99.20 | 12.00 |
|  | Wtdbg2 | 157.75 | 2.68/14 | 1.87/19 | 326 | 19.78 | 94.80 | 2.10 |
| *D. melanogaster* (62X) | NextDenovo | 134.34 | 18.11/4 | 15.68/4 | 196 | 30.99 | 98.70 | 1.07 |
|  | Necat | 144.01 | 19.55/4 | 15.90/4 | 1,200 | 25.86 | 98.70 | 2.45 |
|  | Canu | 154.94 | 8.58/6 | 5.68/7 | 1,738 | 23.53 | 98.80 | 45.55 |
|  | Flye | 135.82 | 18.89/4 | 17.32/4 | 335 | 29.97 | 98.80 | 1.58 |
|  | Wtdbg2 | 137.49 | 6.32/7 | 5.33/9 | 919 | 26.07 | 97.20 | 0.57 |
| *O. sativa* (230X) | NextDenovo | 392.56 | 30.55/6 | 18.00/9 | 81 | 26.45 | 98.60 | 13.05 |
|  | Necat | 394.40 | 25.44/7 | 17.86/9 | 183 | 25.83 | 98.70 | 10.85 |
|  | Canu | 395.23 | 11.57/13 | 9.41/15 | 204 | 24.94 | 98.70 | 728.78 |
|  | Flye | 403.45 | 11.10/14 | 7.84/18 | 115 | 24.76 | 98.70 | 25.02 |
|  | Wtdbg2 | 488.33 | 0.96/88 | 0.81/95 | 553 | 17.90 | 94.10 | 5.85 |
| *Z. mays* (51X) | NextDenovo | 2,118.82 | 44.44/17 | 37.90/21 | 700 | 20.74 | 98.20 | 75.90 |
|  | Necat | 2,171.54 | 22.76/32 | 17.71/38 | 3,307 | 20.41 | 98.20 | 87.87 |
|  | Canu | 2,240.87 | 0.65/950 | 0.62/995 | 6,284 | 19.14 | 98.10 | 1,741.77 |
|  | Flye | 2,122.73 | 2.87/222 | 2.59/242 | 863 | 20.63 | 98.20 | - |
|  | Wtdbg2 | 4,068.86 | 0.07/11298 | 0.05/13848 | 22,258 | 14.07 | 97.00 | - |

NG50 is the length N that 50% of the reference genome is covered in contigs with length $\geq N$ . LG50 is the number of contigs with length ≥ NG50. NGA50 is NG50 of aligned blocks that are obtained by breaking contigs at misassembly events and removing all unaligned bases. LGA50 is the number of aligned blocks with length ≥ NGA50. Misassemblies and QV are evaluated by QUAST, where QV is defined as $-10 \times \log_{10}(\frac{\# \text{ mismatches per } 100 \text{ kbp} + \# \text{ indels per } 100 \text{ kbp}}{100 \text{ kbp}})$ . Gene completeness is
represented by the complete BUSCO values. QV and gene completeness were evaluated using the polished assemblies. The genomes of *A. thaliana*, *D. melanogaster* and *O. sativa* were assembled on the same computer with 60 CPUs and 504 GB RAM of memory. The *Z. mays* genome, assembled by NextDenovo, Necat and Canu, was run on a computer cluster with 7 nodes each with 32 CPUs and 256 GB RAM memory, and assembled by Fly and Wtdbg2 run on a fat computer node.

In terms of running time, for the small (*D. melanogaster* and *A. thaliana*) or medium sized genomes (*O. sativa*), NextDenovo is faster than Canu and Flye. For the repeat-rich *Z. mays* genome, NextDenovo was 23 times faster than Canu, and slightly faster than Necat, but slower than Flye owing to the limitations of the CTA algorithm. Wtdbg2 was the fastest for all genomes. It should be noted that time consumption may vary when different parameters are used. In addition, NextDenovo can distribute almost all subtasks to run on different computer nodes in parallel, and a subtask typically only required 32~64 GB peak memory, so for most genomes NextDenovo can complete genome assembly in a day, when running on dozens of computer nodes.

### Assembly of 35 human genomes by NextDenovo and comparative analysis of segmental duplications between humans

We envisage that the NextDenovo program will potentiate population-scale long-read assemblies, which will in turn facilitate the construction of human pan-genomes using Nanopore long read sequencing at low cost. Here, we collected blood samples from 35 humans with diverse ethnicities, including 13 from Africa, six from East Asia, four from Southeast Asia, six from South Asia, two from the Middle East, two from Europe, one from Oceania, and one from America (**Fig. 2a, Supplementary Table 3**). Principal component analysis (PCA) based on single nucleotide polymorphisms (SNPs) with integration of the 1000 Genomes Project dataset indicated that the 35 genomes together covered much of the genetic diversity present in modern humans (**Supplementary Fig. 1**). For each individual, >150 Gb long reads (mean length 21 kb) were sequenced using the Oxford Nanopore long-read sequencing platform. Each individual contained approximately 12,615 (~0.49× in coverage) ultra-long reads (>100 kb), which enabled contiguous assembly of complex regions in the human genome^8, 10, 21, 22^. In addition, for each individual, ~150 Gb of short reads (100 bp) were sequenced for error polishing and correction.

**Fig. 2:**
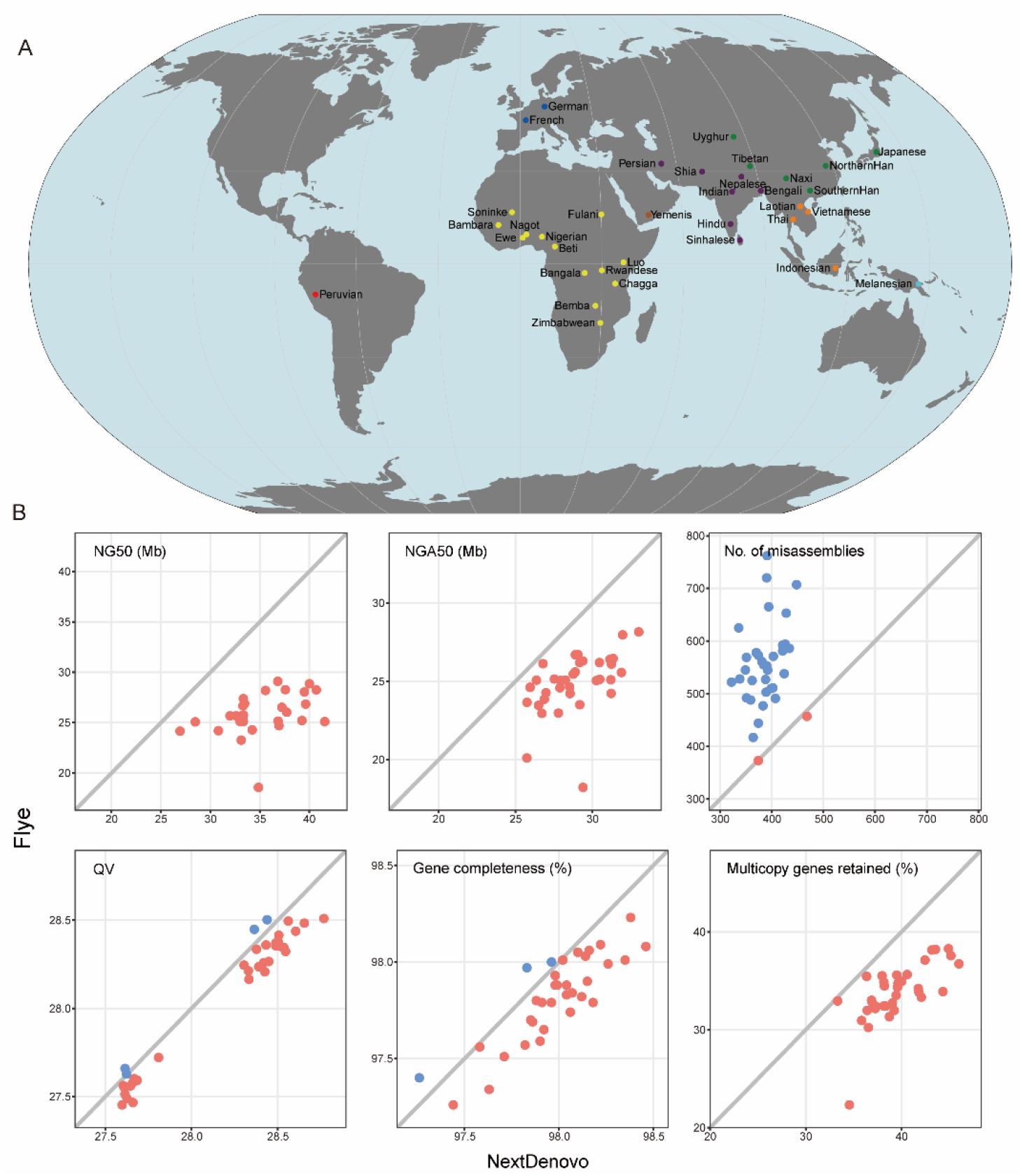
*De novo* assembly of 35 human genomes. (A) Geographical location of the 35 individuals sequenced. (B) Comparison of 35 human assemblies between NextDenovo and Flye. NG50 is the length N such that 50% of the reference genome is covered in contigs with length ≥ N. LG50 is the number of contigs with length ≥ NG50. NGA50 is NG50 of the aligned blocks that are obtained by breaking contigs at misassembly events and removing all unaligned bases. LGA50 is the number of aligned blocks with length ? NGA50. Misassemblies and QV were evaluated by QUAST, where QV is defined as 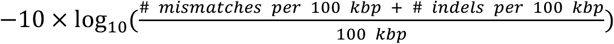. Gene completeness and “multicopy genes retained” are reported by asmgene, “multicopy genes retained” corresponds to the percentage of multicopy genes in the reference genome that remain multicopy genes in the assembly. QV, Gene completeness and “multicopy genes retained” were evaluated using the polished assemblies. The metrics represented by the red points are larger than the metrics represented by the blue points.

We firstly evaluated the performance of NextDenovo as compared with Flye in relation to human genome assembly (**Fig. 2b**, **Supplementary Table 4**). On average, NextDenovo and Flye produced similar assembly sizes (2.83 Gb) with about 90.84% genome coverage, but the assemblies produced by NextDenovo covered more single-copy genes (97.99% *vs*. 97.82%) and retained more multi-copy genes (39.60% *vs*. 33.93%) than the Flye assemblies (**Supplementary Table 4**). Moreover, as with the results of the maize and rice genome assemblies, the NextDenovo assemblies contained longer (1.03-1.61-fold larger NGA50) and fewer contigs (68.18%-96.97% of LGA50) than the Flye assemblies for all 35 genomes. More importantly, the NextDenovo assemblies contained 388 misassemblies on average, ~70% of that of the Flye assemblies, whilst the NextDenovo assemblies also had a slightly larger average QV than the Flye assemblies (28.17 *vs*. 28.06).

Segmental duplications (SDs) are complex segments of DNA with near-identical sequences that are difficult to assemble by short reads; they nevertheless constitute important sources of structural diversity in the human genome and are associated with many human diseases^23, 24^. The use of long-read genome assembly techniques has facilitated the detection of SDs^24, 25^. Here, by using the “Brisk Inference of Segmental duplication Evolutionary structure” (BISER)^26^, we identified an average of 133.6 Mbp of non-redundant SD sequences per individual (**Supplementary Table 5**), corresponding to ~4.7% of the human genome. Our results showed a notable correlation between total SD size and genome size (R2 = 0.9641, p < 2.2e-16, **Supplementary Fig. 2**). We further identified African-specific SD hotspots, based on the difference of SD frequency between African and non-African assemblies (see Methods section). Our results showed that the highly differentiated hotspots were enriched in the pericentromeric regions (**Fig.3**), which concurs with the predicted hotspots of genomic instability noted in T2T-CHM13^24^.

**Fig. 3:**
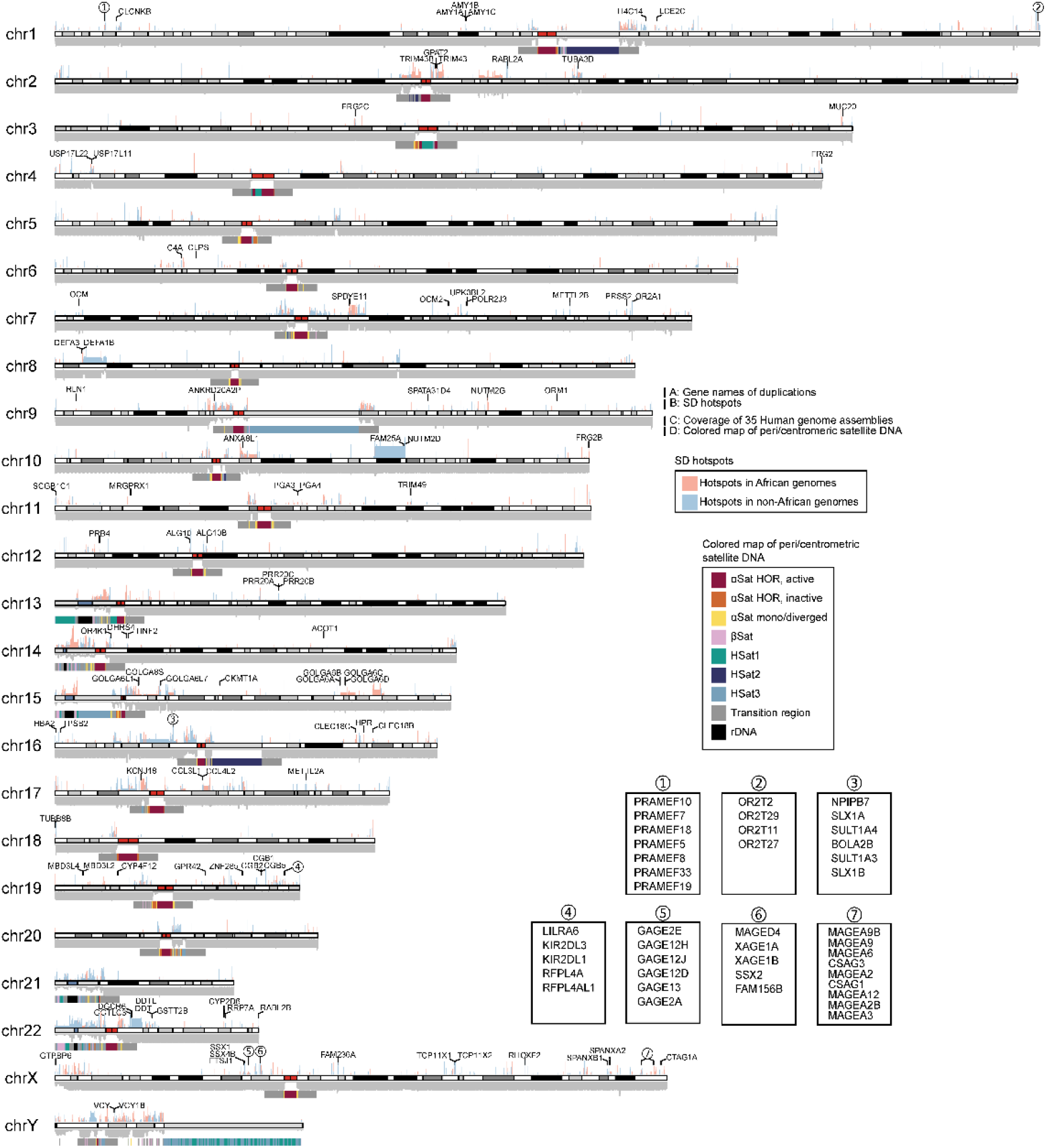
Distribution of duplicate genes and SD hotspots. (A) Gene symbols within duplications (gene names are marked by numbers and are shown in the subfigures). (B) Bar-plots of SD hotspots in African/non-African genomes. (C) Coverage plot of 35 human genome assemblies. (D) Colored map of peri/centromeric satellite DNA (αSat: alpha satellite DNA, βSat: beta satellite DNA, HSat: Human satellite DNA. See^34^ for more detailed definitions). Ideogram plot was built from the T2T-CHM13 (v2) genome. Annotation of peri/centromeric and cytoband regions were downloaded from UCSC (https://hgdownload.soe.ucsc.edu/gbdb/hs1/).

Long-read assembly holds out the promise for the comprehensive discovery of segmental duplications, especially the duplicated genes involved in SDs^24, 25^. We reasoned that these high-quality assemblies should facilitate the detection of gene duplications (**Fig. 3** and **Supplementary Table 6**). In particular, we identified gains of salivary amylase (*AMY1*) gene copies with open reading frames and multiple exons in ten individuals (including 8 Asians and 2 Africans). For example, two individuals from Vietnam and Thailand respectively acquired 4 and 3 additional *AMY1* genes, which may have served to improve their ability to digest starchy foods such as rice. Indeed, the acquisition of additional copies of the *AMY1* gene is known to be a characteristic of populations with a high-starch diet^27^, especially East and South East Asians. Additionally, four clusters of gene families, including preferentially expressed antigen of melanoma (PRAME), olfactory receptor (OR), G antigen (GAGE) and melanoma-associated antigen (MAGEA), exhibited dense clusters of SDs with paralogous genes (**Fig. 3**). Therefore, long-read sequencing makes it possible to accurately assemble those genomic regions that are characterized by highly similar paralogous clusters, including those containing expanded tandemly duplicated genes.

## Discussion

NextDenovo is not only an accurate error-correction tool, but also an efficient *de novo* assembler, specifically developed for noisy long reads using the CTA strategy. In our evaluation, NextDenovo can correct reads at a faster speed and generate more accurate corrected reads than Canu and Necat. The corrected reads usually have a similar accuracy to the HiFi reads while maintaining the contiguity of raw reads. For the assembly, NextDenovo is much faster than the widely used CTA assembler, Canu. It is at least as fast or even faster than Necat based on different input data. For the small and medium sized genomes, it achieved a faster speed than Flye, but NextDenovo was usually slower than other ATC-based tools for repeat-rich large genomes, because of the additional time-consuming error-correction step. However, on the other hand, with the high accuracy imparted by this error-correction step, NextDenovo can generate more continuous assemblies containing fewer misassemblies. This is particularly true when assembling ONT “ultra-long” reads, since NextDenovo can generate partial or near chromosome-level assemblies, and this applies not only to human genome assembly but also to the assembly of complex plant genomes. Indeed, NextDenovo has been successfully applied to large genome assemblies several times, such as with the ~10.5 Gb *Cycas panzhihuaensis* genome (contigs N50 = 12 Mb)^28^, the ~10.76 Gb allohexaploid oat genome (contig N50 = 75.27 Mb)^29^, the ~40 Gb African lungfish genome (contig N50 = 1.60 Mb)^30^ and the ~48Gb Antarctic krill genome (contig N50 = 178.99kb)^31^. Using ONT “ultra-long” reads, NextDenovo can generate partial or near chromosome-level assemblies. Thus, for the ~4.59 Gb papaver genome^32^, NextDenovo produced an assembly with a contig N50 of 65.57 Mb, the longest length being 178.776 Mb using ~19X ONT “ultra-long” reads and ~86X ONT regular reads. In similar vein, for the 3.69 Gb watermelon genome^33^, NextDenovo produced an assembly in which the 11 longest contigs representing 11 chromosomes using ~57X ONT “ultra-long” reads. Finally, for the ~10.76 Gb allohexaploid oat genome^29^, NextDenovo produced an assembly with a contig N50 of 75.27 Mb, the longest length being 313.87Mb using ~100X ONT “ultra-long” reads.

Currently, because of sequencing errors, NextDenovo cannot be used for haplotype-resolved *de novo* genome assembly without trio binning, although it can detect the LSRs caused by heterozygosity, which is the advantage of assembly with HiFi data. However, ONT is gradually updating new base calling models and chemistries that can improve raw data accuracy, which should eventually make it possible for NextDenovo to perform haplotype-resolved assembly.

## Online Methods

### Alignment and filtering

NextDenovo extracts the ~45X longest reads as seeds, and performs pairwise reads overlapping all input reads and seeds using minimap2^35^. For each seed, NextDenovo partitions it into windows of 64bp and calculates the overlapping depth in windows. A window was defined as a repeat window if it had a depth larger than twice the average depth. A window was defined as a chimeric window if it had a depth less than 3. NextDenovo filters out an alignment if it is completely within the repeat windows and splits this seed if it has chimeric windows.

### LSR detection

NextDenovo first uses the kmer score chain (KSC) algorithm to perform the initial rough correction. The KSC algorithm is adapted from the Falconsense algorithm; it calculates a confidence score using the following formula:

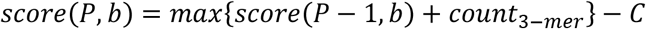

where *C* represents the valid depth at position *P, b* ∈ {*A, T, G, C*, –}, and then determines the correct path using a traceback procedure which starts at the last position *P*. Meanwhile, it records the low-quality positions where the chosen alleles account for ≤ 50% of the total. For each low-quality position, NextDenovo extends it on both sides until there are ≥ 16 consecutive non-low-quality positions. This extended region is defined as a low score region (LSR) if it contains ≥ 4 low-quality positions.

### LSR correction

For a LSR *R* from a seed *S*, all subsequence *Bs* that span this LSR from the overlapping reads of *S* are collected, and a kmer set (*K* = 8) at the 40 bp flanking sequences of *R* is produced. Then for each *B*, the overlapping kmers count between the kmer set from *B* and *R* is calculated as its matched kmer score. NextDenovo sorts all kmer scores of *B*s from large to small and removes all *B*s with a kmer score ≤ *C*, where *C* is smaller than half of its previous kmer score. For the KSC algorithm, deletion errors in the reference sequence are more harmful than insertion errors because the overlapping reads in the regions with insertion errors are not aligned. NextDenovo uses a greedy POA consensus algorithm that adopts a greedy strategy to insert bases in the consensus step to generate a pseudo-LSR seed by using the largest six *B*s ranked by kmer score. All pseudo-LSR seeds from *S* are linked to a long pseudo seed *L*, and all *B*s from *S* are mapped to *L*, and the KSC algorithm is applied to produce a corrected pseudo seed *P*. This procedure is called twice in order to improve the accuracy of the LSRs.

### Graph construction and cleaning

NextDenovo uses two rounds of pairwise overlapping to identify dovetail alignments using a modified Minimap2 between corrected seeds. The first round uses a large batch size and a large repetitive minimizer filtering thread to rapidly detect candidate dovetail alignments. Then for each candidate dovetail alignment, Minimap2 is used again with a smaller repetitive minimizer filtering thread to produce more accurate alignments. Next, a directed string graph is constructed and transitive edges are removed. NextDenovo calculates the average indegree *I* and outdegree *O* of all nodes, and clusters nodes into two categories, repeat nodes and non-repeat nodes. The repeat nodes are defined as nodes with indegree ≥ 1.5*I* or outdegree ≥ 1.5*O*, whereas other nodes are defined as non-repeat nodes. For the paths comprising only non-repeat nodes, the “best overlap graph” (BOG) algorithm is used to remove ambiguous edges. For repeat nodes, NextDenovo first calculates the maximum overlapping identity *I* and maximum overlapping length *L* and maximum transitive score *S* (for an edge *E* from *a* to *c*, if there is node *b*, and there is an edge from *a* to *b* and an edge from *b* to *c*, then the count of *b* is defined as the transitive score of *E*) of out-edges or in-edges, and then removes any edges with overlapping identity <= *i* x *I* and overlapping length *l* x *L* and transitive score 0.5 x *S* (here *i* and *l* are parameters). Subsequently, tips are removed and bubbles are resolved. Finally, for the complex subgraphs which usually contain many repeat nodes connected by only one in-node and one or more out-nodes, or one or more in-nodes and only one out-node, NextDenovo use a series of gradually increasing overlapping identity, overlapping length and transitive score thresholds to remove edges while maintaining connectivity between in-nodes and out-nodes.

### Evaluating error correction

To evaluate the performance of NextDenovo error correction, we simulated about 62X ONT data with N50 length of 20.77 kb from chromosome 1 of the GRCh38 genome using NanoSim (v2.6.0)^36^ and randomly extracted about 72X ONT data with N50 length of 56.77 kb from the chromosome 1 of the CHM13 genome (**Supplementary Table 1**). We next ran NextDenovo, Canu and Necat with the same minimum read lengths to ensure consistency. Finally, we used minimap2 (-x map-ont) to map the corrected data to the reference and assessed their accuracy.

### Evaluating assemblies

We used QUAST for assembly evaluation. For the *A. thaliana, D. melanogaster* and *Z. mays* datasets, we used appropriate NCBI assemblies as the reference genome. For the *O. sativa* dataset, we used the assembly of HiFi data from the same individual by hifiasm (v0.16.1)^4^ as the reference genome. For the human datasets, we used the T2T assembly of CHM13 as the reference genome. The assemblies were further polished with NextPolish using short and long reads and these were subsequently used to evaluate QV and gene completeness. Gene completeness was evaluated with BUSCO for the *A. thaliana*, *D. melanogaster, O.sativa* and *Z. mays* assemblies and paftools (v2.24) asmgene function^4^ for the human assemblies. The commands and parameters used in this study are provided in the Supplementary Information file.

### Human samples for genome sequencing

Sample collection, data release, and paper submission strictly followed all laws of the Ministry of Science and Technology of China (ID: 2021BAT3787). The samples and genome sequence data were not transferred or released outside of China. This study was approved by the Kunming Institute of Zoology Animal Care and Ethics Committee (SMKX-20180715-154) in August 2018. Peripheral blood samples (~5 mL) were collected from people living in China after they provided signed informed consent. Samples were collected from August to October 2018 and were only used in this study.

### DNA extraction, library preparation and sequencing by Nanopore

High-quality genomic DNA was extracted using the SDS (sodium dodecylbenzene sulfonate) method followed by purification with a QIAGEN^®^ Genomic kit (Cat#13343, QIAGEN) according to the standard procedures provided by the manufacturer. DNA degradation and contamination of the extracted DNA were monitored on 1% agarose gels. DNA purity was then detected using a NanoDrop™ One UV-Vis Spectrophotometer (Thermo Fisher Scientific, USA), with OD260/280 ranging from 1.8 to 2.0 and OD260/230 ranging from 2.0 to 2.2. Lastly, DNA concentration was measured using a Qubit^®^ 3.0 Fluorometer (Invitrogen, USA).

In total, 2 μg DNA per sample was used as input material for the ONT (Oxford Nanopore Technologies) library preparations. After the DNA quality was controlled, size-selection of long DNA fragments was performed using the BluePippin system (Sage Science, USA). The DNA fragments were then end repaired, and an A-ligation reaction was conducted using a NEBNext Ultra II End Repair/dA-tailing Kit (Cat# E7546). The adapter in a LSK109 kit was used for further ligation and the Qubit^®^ 3.0 Fluorometer (Invitrogen, USA) was used to quantify the size of the library fragments. Sequencing was then performed on a Nanopore PromethION sequencer (Oxford Nanopore Technologies, UK) at Grandomics Biosciences Co. (Wuhan, China). The Nanopore sequencer output FAST5 files containing signal data and base calling were converted to FAST5 files in FASTQ format with Guppy. The raw reads in fastq format with mean_qscore_template < 7 were then filtered, resulting in pass reads.

### Library preparation and sequencing by MGISEQ2000

Genomic DNA (1 μg) was randomly fragmented by Covaris. The fragmented DNA was selected by an Agencourt AMPure XP-Medium Kit to an average size of 200–400 bp. The selected fragments were subjected to end repair, 3’ adenylation, adaptor ligation, and polymerase chain reaction (PCR) amplification, with the products then being recovered using an AxyPrep Mag PCR Clean-up Kit. The double-stranded PCR products were heat denatured and circularized by the splint oligo sequence. Single-stranded circular DNA (ssCir DNA) was formatted as the final library, and quality controlled. The quality controlled libraries were sequenced on the MGISEQ2000 platform.

### Diversity of 35 human genomes

To determine the diversity of 35 human genomes, we mapped the short reads to the GRCh38 reference assembly using the BWA-MEM (v0.7.15) algorithm^37^. After sorting the reads by coordinates, and removing duplicate reads using SAMtools (v1.8)^38^, HaplotypeCaller and CombineGVCFs in the Genome Analysis Toolkit (GATK, v4.0.4.0)^39^ were used for calling and combining the GVCF files. We then applied the GenotypeGVCFs method in GATK to genotype SNPs based on genome positions from the 1 000 Genomes Project dataset (ftp://ftp.1000genomes.ebi.ac.uk/vol1/ftp/data_collections/1000_genomes_project/release/201_90312_biallelic_SNV_and_INDEL/)^40^. After SNP filtering with “QUAL < 50”, we merged SNPs with the 1000 Genomes Project dataset for principal component analysis.

### Gene, gene duplications and repeat annotations

Gene annotations of the 35 human genomes were performed by mapping GENCODE (v35) annotations^41^ from GRCh38 using Liftoff (v1.6.3)^42^ with the following settings: liftoff - flank 0.1 -sc 0.85 -copies. Duplicate genes were identified based on the following criteria: (1) extra copy number > 1, (2) number of exons > 1, (3) CDS length > 200bp, and (4) containing complete open reading frames (ORF). Repeat annotations were conducted with RepeatMasker (v4.1.3)^43^ and Tandem Repeats Finder (TRF)^44^ RepeatMasker was run with default settings and TRF was run with “trf 2 7 7 80 10 50 15 -l 25 -h -ngs” parameters.

### Segmental duplication (SD) analysis

SDs were identified using BISER (v1.2.3)^26^ based on the soft-masked human genomes. Low-quality SDs were filtered out using the following criteria: (1) <1 kbp in length; (2) >70% overlapping with satellite sequence or > 10% overlapping with simple repeats annotated with RepeatMasker; (3) <90% identical by gap-compressed identity or <50% identical including indels. The pipeline was conducted using a R script (open access on https://github.com/shengwang/35HumanGenome-SDs) and a modified snakemake file download from https://github.com/mrvollger/assembly_workflows/workflows/sedef.smk. Next, we annotated 35 human genomes with unique ancestral units (duplicons) identified by DupMasker^45^. Regions that do not overlap with the duplicons were annotated as new SDs. Finally, we defined the African-specific SD hotspots based on the frequency difference of SDs between African and non-African assemblies. The specific calculation steps were as follows: (1) obtained the non-redundant SD regions for each human assembly, (2) calculated the frequency of SD coverage within African and non-African groups, (3) computed the difference between the frequency of African and non-African of SDs. Regions with a difference much greater than zero were defined as African-specific SD hotspots. We mapped the positional information of SDs from 35 human genome assemblies to the T2T-CHM13 (v2.0, downloaded from https://ftp.ncbi.nlm.nih.gov/genomes/all/GCF/009/914/755/GCF_009914755.1_T2T-CHM13v2.0/GCF_009914755.1_T2T-CHM13v2.0_genomic.fna.gz) genome using the “paftools liftover” tool for visualization. SDs hotspots calculation and visualization were carried out with R packages: tidyverse^46^, rtracklayer^47^, plyranges^48^ and karyoploteR^49^.

## Data availability

The ONT dataset and reference genome for CHM13 were obtained from https://github.com/marbl/CHM13. The ONT, short reads dataset and reference genome for *A. thaliana* were downloaded from BIG Data Center (https://bigd.big.ac.cn/gsa), Beijing Institute of Genomics (BIG), Chinese Academy of Sciences, under accession no. PRJCA005809 (Bioproject), CRR302667 (ONT), CRR302670 (short reads) and GWHBDNP00000001.1 (reference genome). The datasets were obtained from the NCBI Sequence Read Archive: SRR6702603 and SRR6821890 as ONT dataset, SRR6702604 as short reads dataset for *D. melanogaster*, SRR10948639-SRR10948642 as ONT dataset, SRR10948643 as HiFi dataset, SRR10948638 as short reads dataset for *O. sativa*, SRR12482959-SRR12482969 as ONT dataset, SRR11870962 as short reads dataset for *Z. mays*. The reference genomes of *D. melanogaster* and *Z. mays* were downloaded from the NCBI GenBank under accession no. GCA_000001215.4 and GCA_014529475.1, respectively.

All Human data, including raw data and *de novo* assemblies, were deposited at the Genome Sequence Archive (https://ngdc.cncb.ac.cn/gsa-human/) at the National Genomics Data Center, China National Center (NGDC) for Bioinformation/Beijing Institute of Genomics, Chinese Academy of Sciences, under accession number PRJCA006287. Sample collection and data release are permitted by The Ministry of Science and Technology of the People’s Republic of China (permission no. 2021BAT3787). The raw sequencing data of Chinese individuals are available but with restricted access. For more detailed guidance on accessing the data, please refer to the GSA-Human Request Guide for Users (https://ngdc.cncb.ac.cn/gsa-human/document/GSA-Human_Request_Guide_for_Users_us.pdf).

## Code availability

NextDenovo code and benchmarking data are available on https://github.com/Nextomics/NextDenovo and https://nextdenovo.readthedocs.io/en/latest/TEST5.html. The codes and intermediate data of SD analysis are publicly available at https://github.com/shengwang/35HumanGenome-SDs.

## References

1. Eid, J. et al. Real-Time DNA Sequencing from Single Polymerase Molecules. Science 323, 133–138 (2009).

2. Branton, D. et al. The potential and challenges of nanopore sequencing. Nat Biotechnoi 26, 1146–1153 (2008).

3. Wenger, A.M., Peluso, P., Rowell, W.J., Chang, P.C. & Hunkapiller, M.W. Accurate circular consensus long-read sequencing improves variant detection and assembly of a human genome. Nat Biotechnol 37, 155–1162 (2019).

4. Cheng, H., Concepcion, G.T., Feng, X., Zhang, H. & Li, H. Haplotype-resolved de novo assembly using phased assembly graphs with hifiasm. Nat Methods 18, 170–175 (2021).

5. Nurk, S. et al. HiCanu: accurate assembly of segmental duplications, satellites, and allelic variants from high-fidelity long reads. Genome Res 30, 1291–1305 (2020).

6. Lerat, E. Identifying repeats and transposable elements in sequenced genomes: how to find your way through the dense forest of programs. Heredity 104, 520–533 (2010).

7. Jain, M. et al. Nanopore sequencing and assembly of a human genome with ultra-long reads. Nat Biotechnol 36, 338–345 (2018).

8. Nurk, S. et al. The complete sequence of a human genome. Science 376:44–53. (2022).

9. Jain, M. et al. Linear assembly of a human centromere on the Y chromosome. Nat Biotechnol 36, 321–323 (2018).

10. Miga, K.H. et al. Telomere-to-telomere assembly of a complete human X chromosome. Nature 585, 79–84 (2020).

11. Shang, L. et al. A super pan-genomic landscape of rice. Cell Res 32, 878–896 (2022).

12. Tong, X. et al. High-resolution silkworm pan-genome provides genetic insights into artificial selection and ecological adaptation. Nat Commun 13, 5619 (2022).

13. Wang, T. et al. The Human Pangenome Project: a global resource to map genomic diversity. Nature 604, 437–446 (2022).

14. Chen, Y. et al. Efficient assembly of nanopore reads via highly accurate and intact error correction. Nat Commun 12, 60 (2021).

15. Koren, S. et al. Canu: scalable and accurate long-read assembly via adaptive k-mer weighting and repeat separation. Genome Res 27, 722–736 (2017).

16. Ruan, J. & Li, H. Fast and accurate long-read assembly with wtdbg2. Nat Methods 17, 155–158 (2020).

17. Kolmogorov, M., Yuan, J., Lin, Y. & Pevzner, P.A. Assembly of long, error-prone reads using repeat graphs. Nat Biotechnol 37, 540–546 (2019).

18. Hu, J., Fan, J., Sun, Z. & Liu, S. NextPolish: a fast and efficient genome polishing tool for long-read assembly. Bioinformatics 36, 2253–2255 (2019).

19. Lee, C., Grasso, C. & Sharlow, M.F. Multiple sequence alignment using partial order graphs. Bioinformatics 18, 452–464 (2002).

20. Mikheenko, A., Prjibelski, A., Saveliev, V., Antipov, D. & Gurevich, A. Versatile genome assembly evaluation with QUAST-LG. Bioinformatics 34, i142–i150 (2018).

21. Jain, M. et al. Nanopore sequencing and assembly of a human genome with ultra-long reads. Nature Biotechnol 36, 338–345 (2018).

22. Shafin, K. et al. Nanopore sequencing and the Shasta toolkit enable efficient de novo assembly of eleven human genomes. Nat Biotechnol 38, 1044–1053 (2020).

23. Bailey, J.A. & Eichler, E.E. Primate segmental duplications: crucibles of evolution, diversity and disease. Nat Rev Genet 7, 552–564 (2006).

24. Vollger, M.R. et al. Segmental duplications and their variation in a complete human genome. Science 376, eabj6965 (2022).

25. Vollger, M.R. et al. Long-read sequence and assembly of segmental duplications. Nat Methods 16, 88–94 (2019).

26. Išerić, H., Alkan, C., Hach, F. & Numanagić, I. Fast characterization of segmental duplication structure in multiple genome assemblies. Algorithms Mol Biol 17, 1–15 (2022).

27. Perry, G.H. et al. Diet and the evolution of human amylase gene copy number variation. Nat Genet 39, 1256–1260 (2007).

28. Liu, Y. et al. The Cycas genome and the early evolution of seed plants. Nat Plants 8, 389–401 (2022).

29. Peng, Y. et al. Reference genome assemblies reveal the origin and evolution of allohexaploid oat. Nat Genet 54, 1248–1258 (2022).

30. Wang, K. et al. African lungfish genome sheds light on the vertebrate water-to-land transition. Cell 184, 1362–1376 e1318 (2021).

31. Shao, C. et al. The enormous repetitive Antarctic krill genome reveals environmental adaptations and population insights. Cell (2023).

32. Yang, X. et al. Three chromosome-scale Papaver genomes reveal punctuated patchwork evolution of the morphinan and noscapine biosynthesis pathway. Nat Commun 12, 6030 (2021).

33. Deng, Y. et al. A telomere-to-telomere gap-free reference genome of watermelon and its mutation library provide important resources for gene discovery and breeding. Mol Plant 15, 1268–1284 (2022).

34. Altemose, N. et al. Complete genomic and epigenetic maps of human centromeres. Science 376, eabl4178 (2022).

35. Li, H. Minimap2: pairwise alignment for nucleotide sequences. Bioinformatics 34, 3094–3100 (2018).

36. Yang, C., Chu, J., Warren, R.L. & Birol, I. NanoSim: nanopore sequence read simulator based on statistical characterization. GigaScience 6, 1–6 (2017).

37. Li, H. Aligning sequence reads, clone sequences and assembly contigs with BWA-MEM. arXiv e-prints (2013).

38. Li, H. et al. The Sequence Alignment/Map format and SAMtools. Bioinformatics 25, 2078–2079 (2009).

39. McKenna, A. et al. The Genome Analysis Toolkit: a MapReduce framework for analyzing next-generation DNA sequencing data. Genome Res 20, 1297–1303 (2010).

40. Auton, A. et al. A global reference for human genetic variation. Nature 526, 68–74 (2015).

41. Harrow, J. et al. GENCODE: The reference human genome annotation for The ENCODE Project. Genome Res 22, 1760–1774 (2012).

42. Shumate, A. & Salzberg, S.L. Liftoff: accurate mapping of gene annotations. Bioinformatics 37, 1639–1643 (2021).

43. Smit, A., Hubley, R. & Green, P. RepeatMasker Open-4.0. 2013-2015 289–300. Available online at http://www.repeatmasker.org (accessed March 18, 2020) (2015).

44. Benson, G. Tandem repeats finder: a program to analyze DNA sequences. Nucleic Acids Res 27, 573–580 (1999).

45. Jiang, Z., Hubley, R., Smit, A. & Eichler, E.E. DupMasker: A tool for annotating primate segmental duplications. Genome Res 18, 1362–1368 (2008).

46. Wickham, H. et al. Welcome to the Tidyverse. J Open Source Softw 4, 1686–1686 (2019).

47. Lawrence, M., Gentleman, R. & Carey, V. rtracklayer: an R package for interfacing with genome browsers. Bioinformatics 25, 1841–1842 (2009).

48. Lee, S., Cook, D. & Lawrence, M. plyranges: a grammar of genomic data transformation. Genome Biol 20, 4 (2019).

49. Gel, B. & Serra, E. karyoploteR: an R/Bioconductor package to plot customizable genomes displaying arbitrary data. Bioinformatics 33, 3088–3090 (2017).

